# *Drosophila* adult muscle precursor cells contribute to motor axon pathfinding and proper innervation of embryonic muscles

**DOI:** 10.1101/604215

**Authors:** Guillaume Lavergne, Krzysztof Jagla

## Abstract

Adult Muscle Precursors (AMPs), the *Drosophila* muscle stem cells, arise from the asymmetric cell divisions of a subset of muscle progenitors ([1]) and are characterized by the persistent expression of the myogenic transcription factor Twist ([2]) and activation of the Notch pathway ([3]; [4]). They occupy stereotyped positions in the vicinity of developing body wall muscles, stay quiescent and undifferentiated during embryonic life and are reactivated during second larval instar ([5]; [6]) to generate muscles of the adult fly. Strikingly, AMPs are also located in the path of intersegmental (ISN) and segmental (SN) motor neuron branches ([2]; [7]). However, their role and interactions with the motor neurons have not yet been analyzed in details. Here, using AMP sensor line revealing cell membrane extensions we show that the navigating ISN first contacts the dorso-lateral (DL-AMPs) and then the dorsal AMP (D-AMP) marking the end of its trajectory. In parallel, the segmental nerve SNa innervating lateral muscles targets the lateral AMPs (L-AMPs). *In vivo* analyses of AMPs behavior highlight an active filopodial dynamic of AMPs toward the ISN and SNa suggesting they could guide motor axons and contribute to muscle innervations. Indeed, our data show that loss or mispositioning of L-AMPs affect the SNa motor axons pathfinding and branching, leading to loss or aberrant muscle innervation. The finding of a transient expression of the guidance molecule Sidestep in L-AMPs suggests its implication in this process. Thus, proper muscle innervation does not only rely on the dialogue between the motor neurons and the muscles, but also on the AMP cells. AMPs represent spatial landmarks for navigating motor neurons and their positioning is critical for the muscles innervation in the lateral region.

## Results and discussion

### AMPs actively interact with the motor axons

During *Drosophila* embryogenesis, we can distinguish a stereotypic pattern of AMPs per abdominal hemisegment in ventral (V-AMP), lateral (L-AMPs), dorsolateral (DL-AMPs) and dorsal (D-AMPs) positions (Fig. 1B). Here we investigated the relationship between AMPs and motor axons and their dynamics during development using embryos carrying the M6-gapGFP transgene, which allows visualization of the membrane of AMP cells ([6]). We found that the intersegmental nerve (ISN) established contacts with the DL-AMPs during the embryonic stage 13 (Fig. 1A-B) and then navigated toward the D-AMP to contact it at stage 15 (Fig. 1C-D). Within the lateral field, the segmental nerve a (SNa) is sub-divided into two branches, dorsal (D-SNa), which innervates the lateral transverse muscles (LTs 1-4), and lateral (L-SNa) targeting the Segmental Border Muscle (SBM). The SNa sub-division takes place during stage 15 and we observed that the L-SNa branch migrated towards the L-AMPs before innervating the SBM (Fig. 1C-D). In parallel, the anterior L-AMP underwent shape changes and directional migration towards the L-SNa (Fig. 1C-C”). In a similar way, one of the DL-AMPs moves dorsally following ISN migration and the D-AMPs appear to extend toward the ISN (Fig. 1C-C””). However, AMPs survival and behavior are not affected in the absence of motor axons as shown in the *prospero* mutant where motor axons fail to exit the CNS (Fig. S1 and [8]). To better characterize dynamics of AMP-motor axons interactions we used the M6-GAL4-driven UAS-Life-actin GFP reporter line that allows *in vivo* visualization of both the motor axons and the AMPs (Fig. 1E-L). Live imaging revealed that among the numerous oriented cytoplasmic projections sent out by the AMPs; those contacting the growth cones of motor axons became stabilized (Fig. 1F,J, L). In particular, stabilization of filopodial connections between L-AMPs and SNa coincided with the setting of the SNa branching point and specification of its lateral branch (Fig. 1I-L). We also found oriented filopodial dynamics in the dorsal region with the contact between D-AMP projections and ISN growth cone prior to ISN migration toward the D-AMP (Fig. 1E-H). As previously demonstrated, muscle founders are needed for terminal defasciculation of the main motor axon branches ([9]). In this context, AMPs positioning and the fact that they actively engage with the navigating motor axons, might also participate in this process by acting as spatial check-points either inducing and or attracting targeted defasciculation of ISN and SNa.

**Figure 1.**
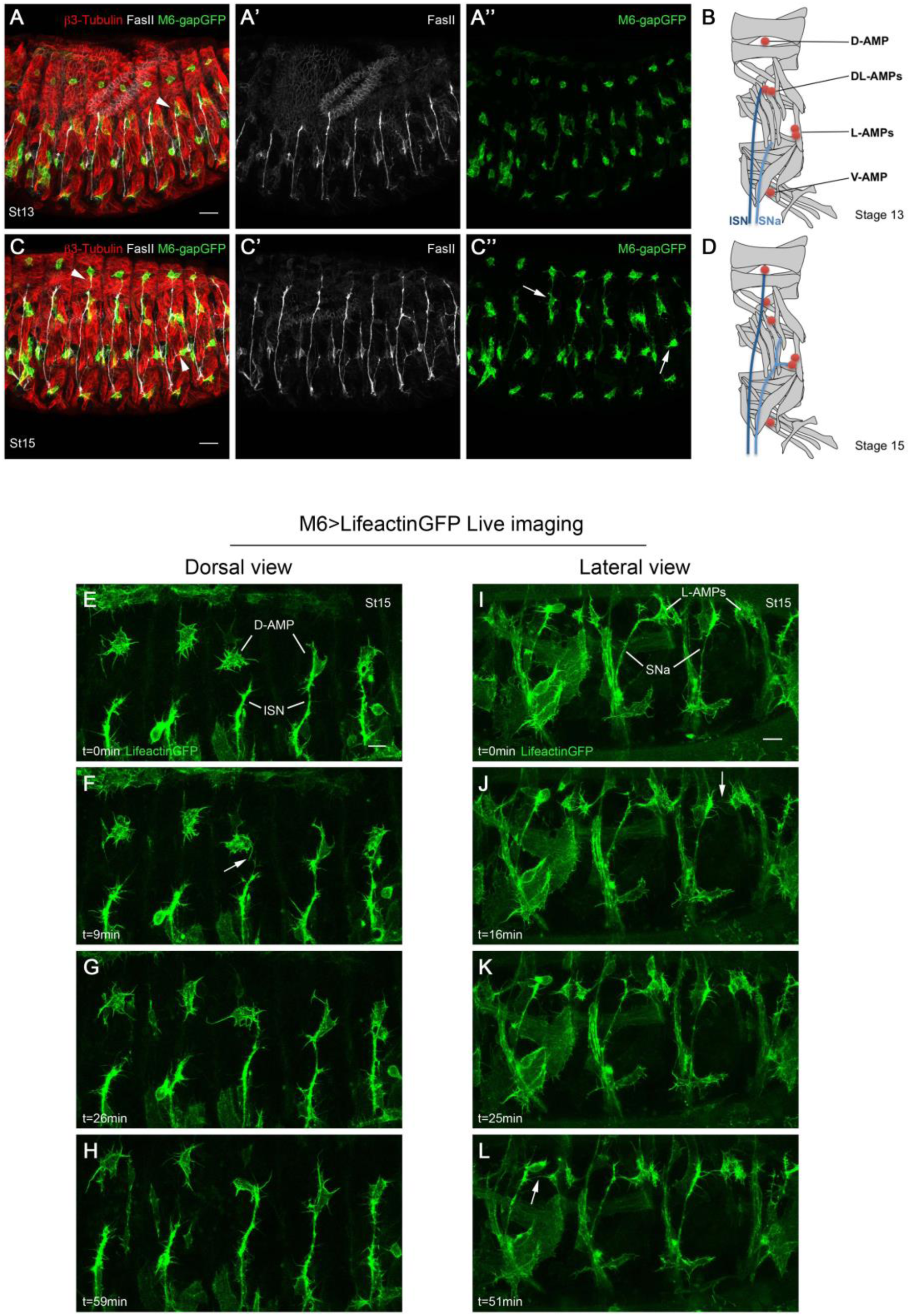
AMPs actively interact with the motor axons. (**A-B’’**) Lateral view of stage 13 (**A-A’’**) or stage 15 (**C-C’’**) M6-gapGFP embryos showing the embryonic pattern of motor axons (stained with anti-Fas2) and AMPs (visualized with M6-GapGFP) associated with muscles (marked by anti-β3-Tubulin). Arrowhead in **A**’ indicates an example of contact between DL-AMPs and the growing ISN. In **C**, arrowheads point toward the contact made between SNa/L-AMPs and ISN/D-AMPs. In C’’, white arrows illustrate the mobility of DL-AMPs and L-AMPs in late embryogenesis. (**B,D**) Scheme representing muscle pattern (gray), the AMPs (red) and the two major nerve branches ISN (dark blue) and SNa (light blue) within an abdominal hemisegment. (**E-L**) Selected time-points from supplemental movies 1 and 2, showing *in vivo* dorsal view (**E-H**) or lateral view (**I-L**) of M6-GAL4>UAS-LifeactinGFP stage 15 embryos. The white arrows mark the first contacts between filopodia from SNa and L-AMPs (**F,L**) or a D-AMP and the ISN (**F**). Scale bars represent 20 µm (**A,C**) or 10 µm (**E,I**)

### L-AMPs are required for the lateral sub-branching of the SNa

To investigate the impact of L-AMPs on the SNa pathway and branching, we first assessed the effect of a genetic ablation of the AMP cells using the M6-GAL4-driven expression of the pro-apoptotic gene *reaper*. This enabled targeted induction of apoptosis selectively in AMPs with occasional AMP cell loss and without noticeable defects in motor neuron networks in which M6-GAL4 is also expressed. This differential effect could be due to a lower expression level of M6-GAL4 in motor neurons than in AMPs. Importantly, in 84% of hemisegments (n=36), complete loss of L-AMPs was associated with absence of the lateral branch of SNa (L-SNa), strongly suggesting that L-AMPs play an instructive role in L-SNa formation and/or stabilization (Fig. 2F-J). In contrast, loss of L-SNa in hemisegments where the L-AMPs were still present occurred in 11% of analyzed hemisegments (n=147). M6-GAL4-induced apoptosis created a context in which loss of the L-SNa branch was observed in L-AMP-devoid segments where the L-SNa target muscle (SBM) was still present (Fig. 2F-J). This suggests that L-SNa branch formation might not be dependent on its muscle target, and so prompted us to test whether the L-SNa would form in an SBM-devoid context. We targeted *reaper* expression to the developing SBM using the previously described SBM-GAL4 driver (gift from A. Michelson). The SBM-GAL4-driven apoptosis resulted in systematic loss of the SBM, with no major impact on the L-AMPs. In this SBM-devoid context but with L-AMP cells correctly located, the L-SNa branch formed normally (Fig. 2K-O). By contrast, in a subset of SBM-deficient embryos L-AMP misplacement toward the navigating SNa resulted in a shortened L-SNa (Fig. 2L). These observations thus suggest an instructive role for AMPs in L-SNa establishment, and reveal that SBM might not be needed for this process and is at least dispensable for its stabilization. To further test the role of L-AMPs in lateral defasciculation of the SNa we analyzed different genetic contexts in which AMP specification is affected. We first induced a perturbation of asymmetric cell divisions. To adversely affect divisions of progenitor cells that give rise to AMPs we ectopically expressed the asymmetry determinant Numb using the pan-mesodermal driver Twist-GAL4 ([10]). In the lateral region, this led predominantly to the loss of one of the L-AMPs and a duplication of the SBM. However, in a small subset of hemisegments (less than 1%) we observed rare loss of both L-AMPs with no impact on SBM (often duplicated) (Fig. 3B-B”). In this context, the L-SNa was absent in 100% of hemisegments (n=10) (Fig. 3B-B”) supporting the view that L-AMPs are required for L-SNa branching. These findings are also consistent with the effects of generalized mesodermal expression of the identity gene *Pox meso*, which can lead to a loss of L-AMPs ([11]), and consequently affects lateral SNa formation in L-AMP-devoid segments (Fig. 3C-C”). Interestingly, pan-mesodermal expression of *Pox meso* can also induce misplacement of L-AMPs along the SBM leading to aberrant L-SNa trajectory (Fig. 3C-C”). Hence L-AMPs and their spatial positioning appear critical to achieve the formation and correct pathfinding of the L-SNa.

**Figure 2.**
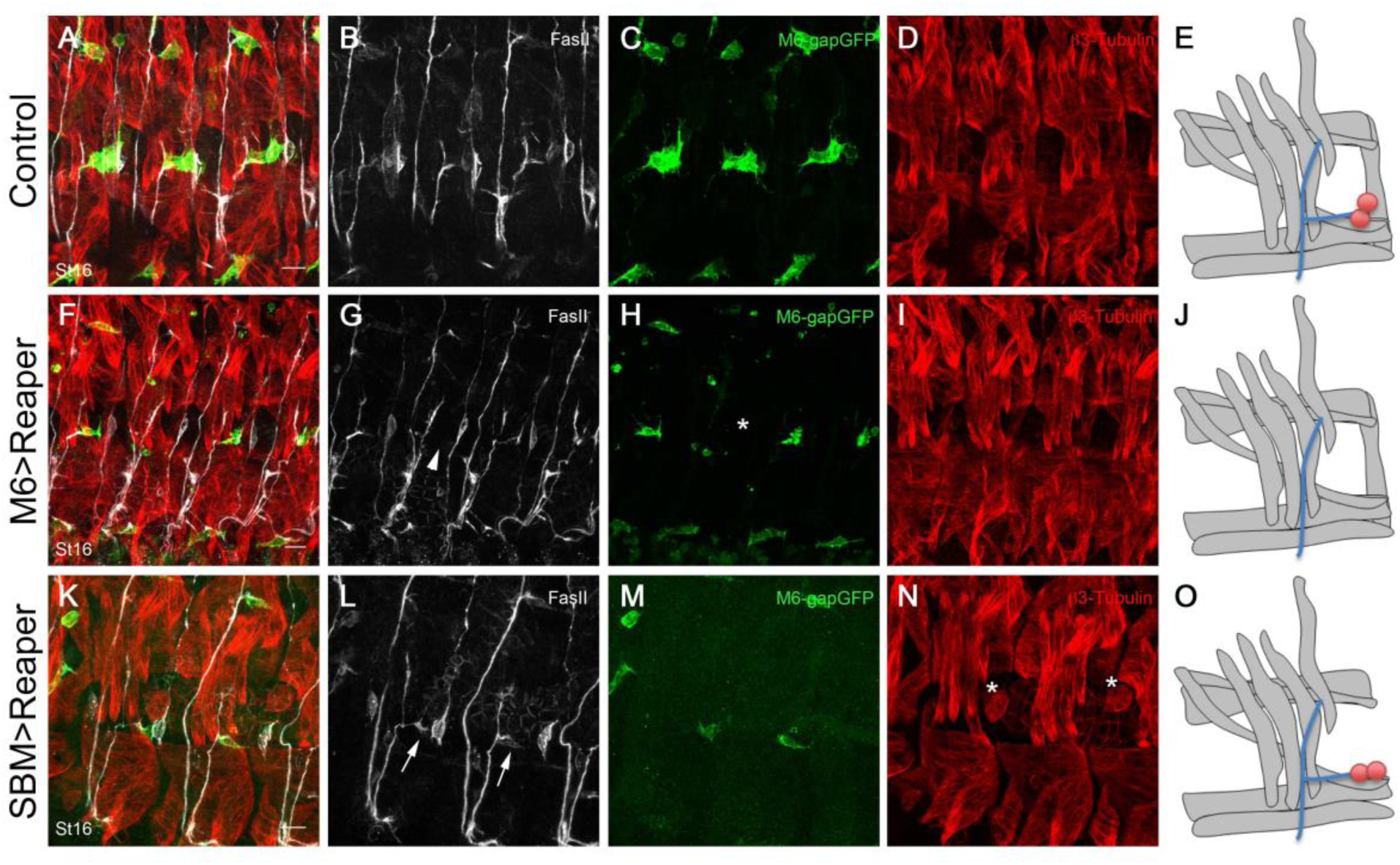
Effects of targeted apoptosis on lateral SNa branching. Zoomed lateral view of a stage 16 M6-gapGFP as a control (**A-D**); M6-gapGFP;SBM-GAL4>UAS-Reaper (**F-I**) and M6-gapGFP;M6-GAL4>UAS-Reaper (**K-N**) embryos. (**F-I**) Complete apoptosis of L-AMPs is observed in one hemisegment marked by a white asterisk. The same hemi-segment also displays a loss of L-SNa (white arrowhead). (**K-N**) Apoptosis of SBM muscle is marked by a white asterisk. Persistence of the L-SNa can be observed (white arrows) in contact with remnant L-AMP even when the SBM is missing. Note that length of the SNa appears dependent of the positioning of the L-AMPs. (**E,J,O**) Results are illustrated by recapitulative schemes with muscles in gray, AMPs in red and SNa in blue. Scale bars represent 10 µm.

**Figure 3.**
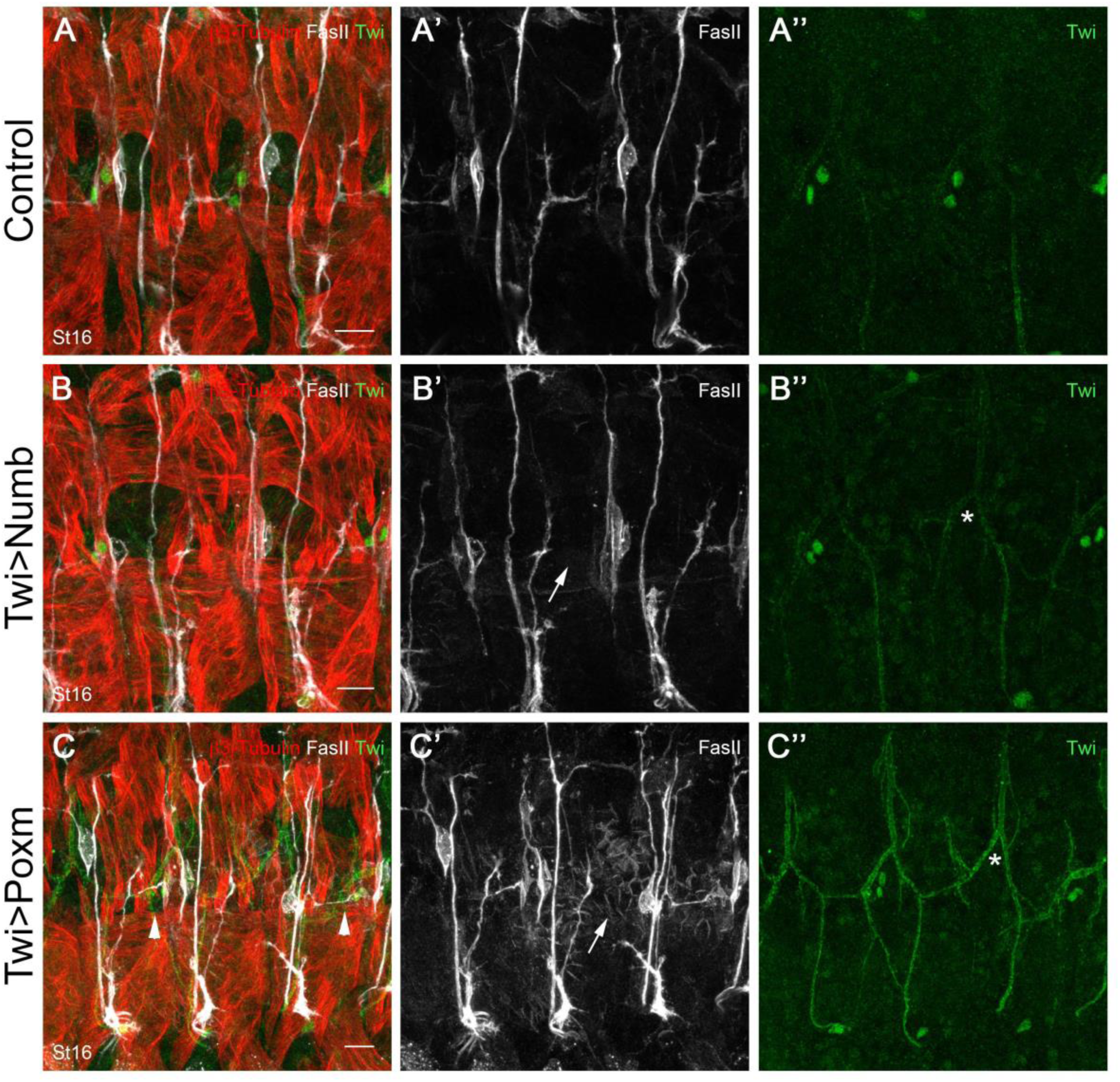
Specification of L-AMPs is required for the formation of the L-SNa. Zoomed lateral view of either Twist-GAL4 (**A-A’’**), Twist-GAL4>UAS-Numb (**B-B’’**) or Twist-GAL4>UAS-Pox meso (**C-C’’**) stage 16 embryos. AMPs are revealed by the staining with the Twist antibody shown in green. Ectopic expression of Numb and Pox meso in the mesoderm is sufficient to affect the specification of AMPs. Absence of L-AMPs in a given hemisegment is marked with white asterisks (**B’’, C’’**). Segments lacking L-AMPs also present absence of L-SNa indicated with a white arrow (**B’,C’**). (**C**) White arrowheads point toward dorsally mispositioned L-AMPs and the resulting L-SNa stabilized at a more dorsal position than in control segments. Scale bars represent 10 µm.

### AMPs express Sidestep, involved in motor axon guidance

The findings described above suggest that L-AMPs are a source of attractive signals that promote lateral sub-branching of the SNa making it competent to innervate the SBM. Interestingly, the SBM is the only lateral muscle innervated by the Connectin-positive SNa, which does not express this homophilic target recognition molecule ([12]; [13]). In such a context, L-AMP-mediated lateral sub-branching of SNa offers a way to drive L-SNa to its specific muscle target. Since L-AMPs seem not to express Connectin either (Fig. S2), in contrary to what had been previously suggested ([14]), their role in attracting SNa and inducing the L-SNa sub-branching might rely on other guidance molecules.

It has been previously shown that the mutants of *sidestep* and *beat-1a*, encoding interacting membrane proteins of the Immunoglobulin superfamily, displayed loss of L-SNa, a phenotype similar to the one observed when L-AMPs are missing ([15]; [16]). However, the mechanisms leading to the loss of the L-SNa in *sidestep* and *beat-1a* mutants have not been elucidated. Also, the embryonic expression pattern of *sidestep* has been only partially characterized ([16]; [17]). We therefore decided to test Sidestep protein distribution at the time when L-SNa sub-branching is taking place. By examining stage 14 to 15 embryos we found a previously unreported faint and transient expression of Sidestep specifically in L-AMPs (Fig. 4A,A’). To confirm this observation we analyzed the expression of Sidestep in a mutant for *beat-1a.* It has been reported ([17]) that the contact of Beat-1a expressing motor axons with Sidestep-expressing cells leads to a negative regulation of the expression of *sidestep*. If this contact is missing, cells normally expressing *sidestep* transiently and at a low level will continue to do so, leading to continuous and higher Sidestep level in these cells. Analyses of *beat-1a* mutants confirmed that the L-AMPs are Sidestep-expressing cells and that Sidestep expression onset coincides with L-SNa sub-branching (Fig. 4B-C’). The high Sidestep expression resulting from the lack of *beat-1a* was still detected in late-stage embryos in which it became gradually restricted to the most anterior L-AMPs (Fig. 4D,D’). This late differential Sidestep expression may point to a leading role for the anterior L-AMPs in the process of interaction with SNa and in its lateral sub-branching. Importantly, the SBM does not express Sidestep, making the L-AMPs the only Sidestep-expressing cells in the L-SNa pathway. Thus, this newly reported expression pattern strongly suggest that L-AMPs attract the L-SNa through the temporally and spatially restricted expression of *sidestep*.

**Figure 4.**
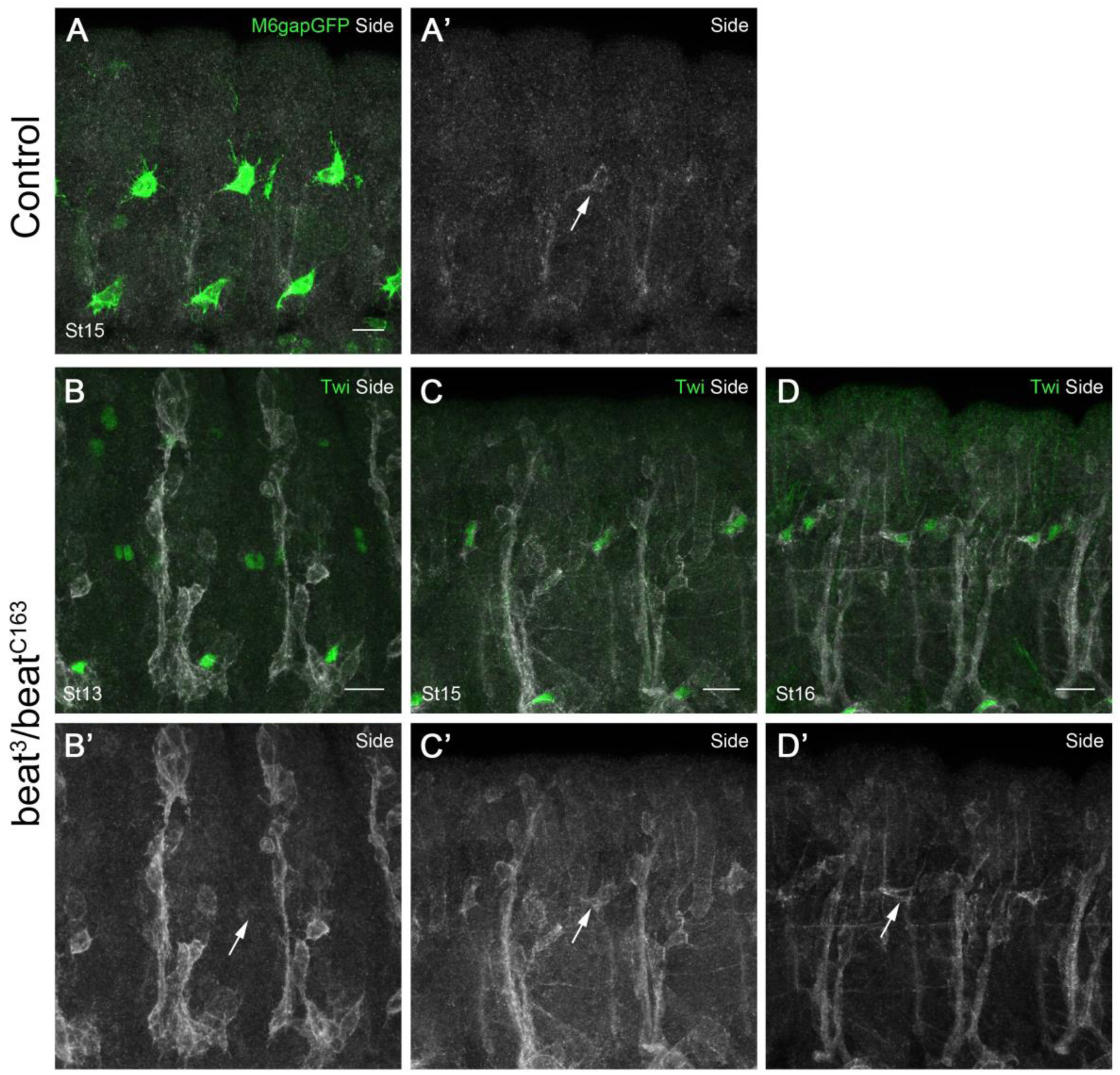
L-AMPs transiently express the guidance molecule Sidestep. (**A,A’**) Zoomed lateral view of a M6-gapGFP stage 15 embryo. Immunostaining with the anti-Sidestep antibody reveals a weak expression in L-AMPs (white arrows). (**B-D’**) In *beat-1a* mutant embryo Sidestep expression appears higher in Sidestep-positive cells. At stage 13 L-AMPs are Sidestep-negative (**B,B’**). Increased Sidestep expression in L-AMPs could be detected at stage 15 (**C,C’**) and persists in the anterior L-AMP in stage 16 *beat-1a* mutant embryos (**D,D’**). Scale bars represent 10 µm.

Interestingly, the *sidestep* mutants also display a stall phenotype of the ISN ([16]) suggestive of a potential role of the D-AMPs. However, we were unable to detect a clear Sidestep expression in D-AMPs in either control or *beat* mutant embryos, indicating that other guiding cues may be in play. In this regard, we could observe that the aberrantly located D-AMPs, after the mesodermal overexpression of the activated form of the Notch receptor (NICD), are able to shift the ISN pathway in their direction, suggesting that they are a source of attracting signals (Fig. S3). Hence both D-AMPs and L-AMPs are able to attract navigating motor axons and influence their trajectories. However, loss of D-AMPs, only reported in pan-mesodermal overexpression of *pox meso*, does not affect the capacity of ISN motor axons to target dorsal muscles (Fig. S3). These results highlight differential requirements of AMPs on motor axons terminal defasciculation, with L-AMPs being mandatory for L-SNa establishment and D-AMPs acting at a guidance level on the ISN.

It is also important to state that the loss of L-SNa in the *sidestep* mutants is not fully penetrant ([16]), suggesting that *sidestep* might not be the only player in L-SNa sub-branching. This could be due to functional redundancy between several members of Side and Beat families comprising 8 and 14 members, respectively ([18]). Expression and function of Side and Beat family members remain largely unexplored, but the fact that Sidestep labels L-AMPs and its paralog, *side VI*, is expressed in the DL-AMPs ([19]) suggests there might be a “Side expression code” that operates in AMPs and makes them competent to interact with navigating motor axons.

Thus, in *Drosophila*, the dynamic interactions and close association between AMPs and the motor axon network contribute to setting ISN trajectory and are required for SNa sub-branching and proper innervation of lateral muscles. This study represents the first demonstration of the role played by muscle stem cells in the establishment of the muscle innervation. Whether motor axons interact with muscle stem cells in vertebrates remains largely unexplored. However, it has recently been shown that muscle stem cells activate and contribute to mouse neuromuscular junction regeneration in response to denervation ([20]) and that depletion of muscle stem cells induced neuromuscular junction degeneration ([21]).

### STAR methods

#### KEY RESOURCES TABLE

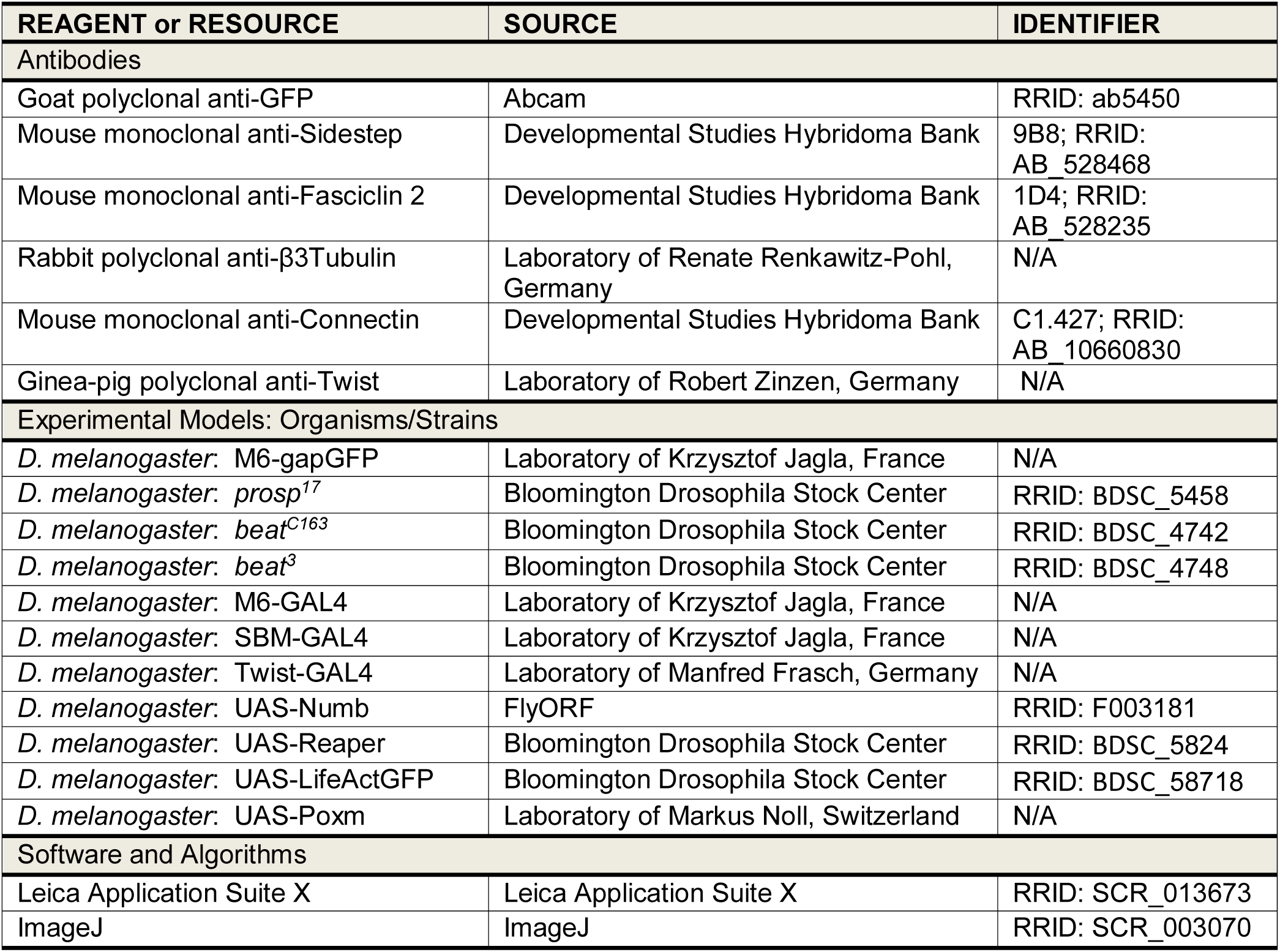

| REAGENT or RESOURCE | SOURCE | IDENTIFIER |
| --- | --- | --- |
| **Antibodies** |  |  |
| Goat polyclonal anti-GFP | Abcam | RRID: ab5450 |
| Mouse monoclonal anti-Sidestep | Developmental Studies Hybridoma Bank | 9B8; RRID: AB_528468 |
| Mouse monoclonal anti-Fasciclin 2 | Developmental Studies Hybridoma Bank | 1D4; RRID: AB_528235 |
| Rabbit polyclonal anti-β3Tubulin | Laboratory of Renate Renkawitz-Pohl, Germany | N/A |
| Mouse monoclonal anti-Connectin | Developmental Studies Hybridoma Bank | C1.427; RRID: AB_10660830 |
| Ginea-pig polyclonal anti-Twist | Laboratory of Robert Zinzen, Germany | N/A |
| **Experimental Models: Organisms/Strains** |  |  |
| *D. melanogaster*: M6-gapGFP | Laboratory of Krzysztof Jagla, France | N/A |
| *D. melanogaster*: *prosp*<sup>17</sup> | Bloomington Drosophila Stock Center | RRID: BDSC_5458 |
| *D. melanogaster*: *beat*<sup>C163</sup> | Bloomington Drosophila Stock Center | RRID: BDSC_4742 |
| *D. melanogaster*: *beat*<sup>3</sup> | Bloomington Drosophila Stock Center | RRID: BDSC_4748 |
| *D. melanogaster*: M6-GAL4 | Laboratory of Krzysztof Jagla, France | N/A |
| *D. melanogaster*: SBM-GAL4 | Laboratory of Krzysztof Jagla, France | N/A |
| *D. melanogaster*: Twist-GAL4 | Laboratory of Manfred Frasch, Germany | N/A |
| *D. melanogaster*: UAS-Numb | FlyORF | RRID: F003181 |
| *D. melanogaster*: UAS-Reaper | Bloomington Drosophila Stock Center | RRID: BDSC_5824 |
| *D. melanogaster*: UAS-LifeActGFP | Bloomington Drosophila Stock Center | RRID: BDSC_58718 |
| *D. melanogaster*: UAS-Poxm | Laboratory of Markus Noll, Switzerland | N/A |
| **Software and Algorithms** |  |  |
| Leica Application Suite X | Leica Application Suite X | RRID: SCR_013673 |
| ImageJ | ImageJ | RRID: SCR_003070 |

## CONTACT FOR REAGENT AND RESOURCE SHARING

Further information and requests for resources and reagents should be directed to and will be fulfilled by the Lead Contact, Krzysztof Jagla.

## EXPERIMENTAL MODEL AND SUBJECT DETAILS

*Drosophila* stocks were maintained at 25°C. The M6-gapGFP ([6]) strain was used as control. Mutant strains used were *prosp*^*17*^(BL5458), *beat*^*C163*^(BL4742) and *beat*^*3*^(BL4748). The targeted expression experiments were performed using the UAS-GAL4 system ([22]) on the following GAL4 and UAS lines: M6-GAL4 (generated in the lab) and SBM-GAL4 (kindly provided by A. Michelson); Twist-GAL4 (kindly provided by M. Frasch); UAS-Poxm (kindly provided by M. Noll); UAS-Numb (FlyORF:F003181), UAS-Reaper (BL5824), UAS-LifeActGFP (BL58718) from Bloomington Stock Center. Mutant strains were balanced with CyO ActinGFP and TM6B KrGFP, homozygous mutant embryos were selected by absence of corresponding GFP staining. All GAL4-UAS crosses were performed at 25°C.

## METHOD Details

### Immunohistochemical staining and live imaging

Staged embryos were dechorionated, fixed, blocked for 1 h at room temperature in 20% horse serum in PBT and then incubated with primary and secondary antibodies according to a standard procedure ([23]). The primary antibodies used were: goat anti-GFP (1:500, Abcam, ab 5450), guinea-pig anti-Twi (1:1000, kindly provided by R. Zinzen), rabbit anti-β3Tubulin (1:2000, kindly provided by R. Renkawitz-Pohl; Friedrich-Alexander University of Erlangen-Nuremberg, Erlangen, Germany); mouse anti-Fas2 (1D4; 1:400), mouse anti-Con (C1.427; 1:50), mouse anti-Side (9B8; 1:50) from Developmental Studies Hybridoma Bank (University of Iowa). Secondary antibodies were: donkey anti-rabbit, donkey anti-guinea pig, donkey anti-goat and donkey anti-mouse (Jackson Immuno Research Laboratories) conjugated to Alexa 488, CY3 or CY5 fluorochromes (dilution 1:300). Labeled embryos were analyzed using a Leica SP8 confocal microscope equipped with a HyD detector, with a 40X objective. Images were processed with ImageJ.

*In vivo* imaging of M6>LifeActGFP embryos was performed from stage 14 to 16 over 2 h. Films were acquired on a Leica SP8 confocal microscope and analyzed using Imaris software (Bitplane).

## Supplemental Information

### Supplemental figure legends

**Figure S1.**
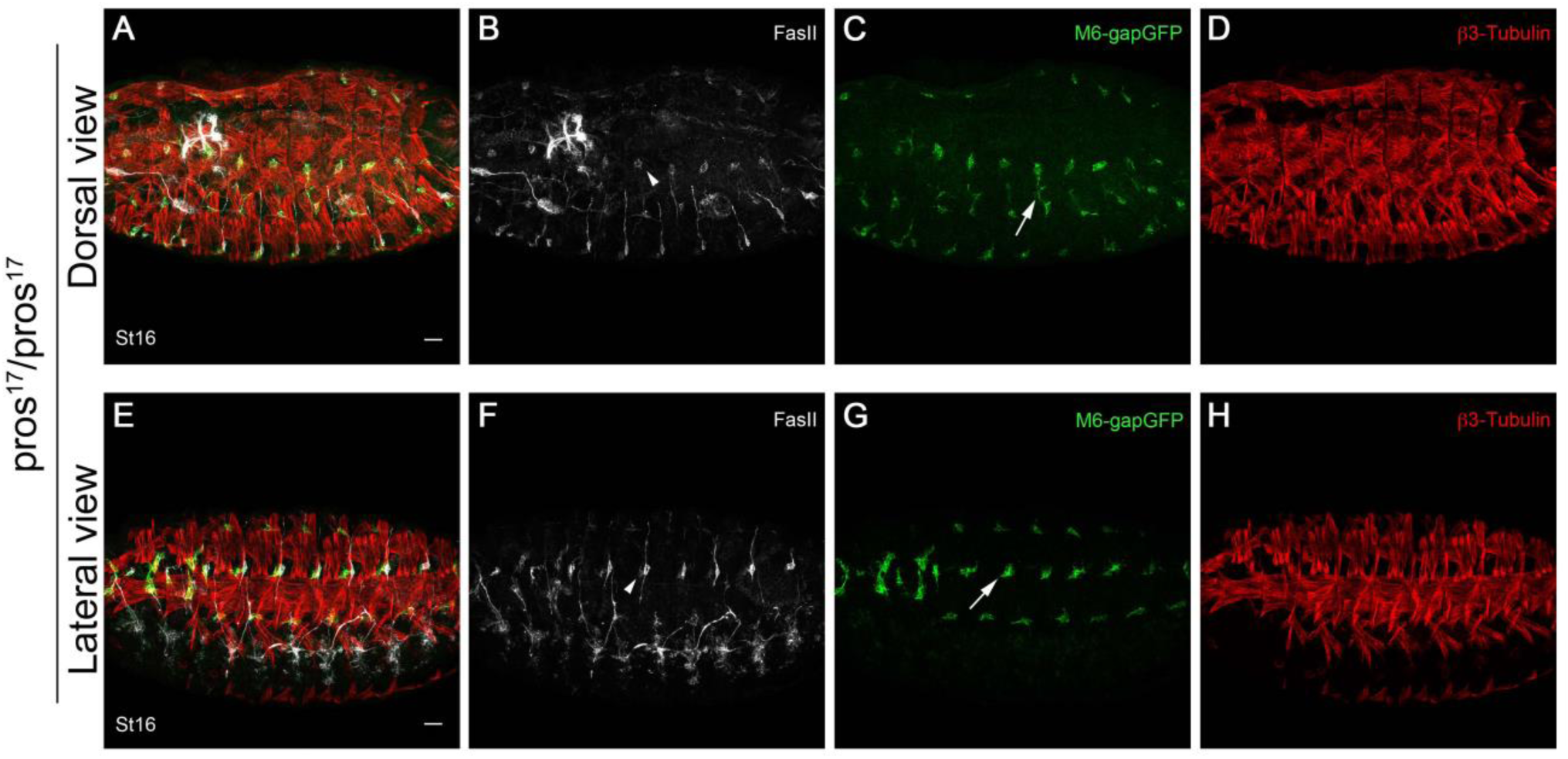
Motor axons are not necessary for AMP behavior. Dorsal view (**A-D**) and lateral view (**E-H**) of stage 16 *prospero* mutants combined with the M6-gapGFP sensor. In this context motor axons do not exit the CNS properly leaving only a minor extension toward the most ventral muscles, and the Fas2 positive extension of the LBD in the dorsal region. White arrows point toward the unaffected migration of DL-AMPs (**C**) and L-AMPs (**G**). Arrowheads highlight the Fas2 expression associated with AMP cells (**B,F**). Scale bars represent 20 µm.

**Figure S2.**
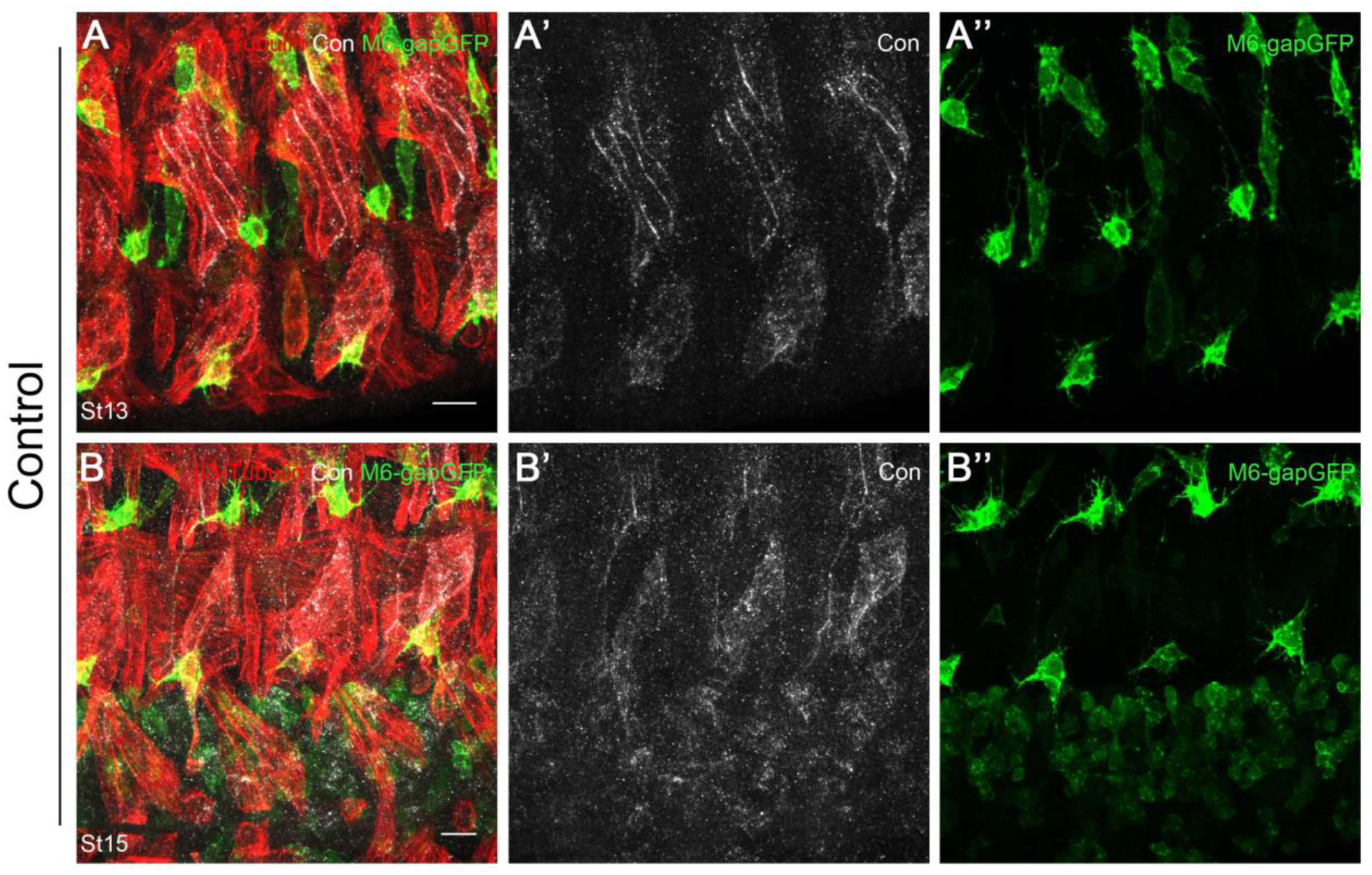
L-AMPs do not express the cell adhesion molecule Connectin. Ventrolateral view of stage 13 (**A-D**) and stage 16 (**E-H**) M6-gapGFP embryos stained with anti-Connectin. Connectin expression is observed in lateral and ventral acute muscles, and in some cells in the CNS. AMPs are Connectin-negative. Scale bars represent 10 µm.

**Figure S3.**
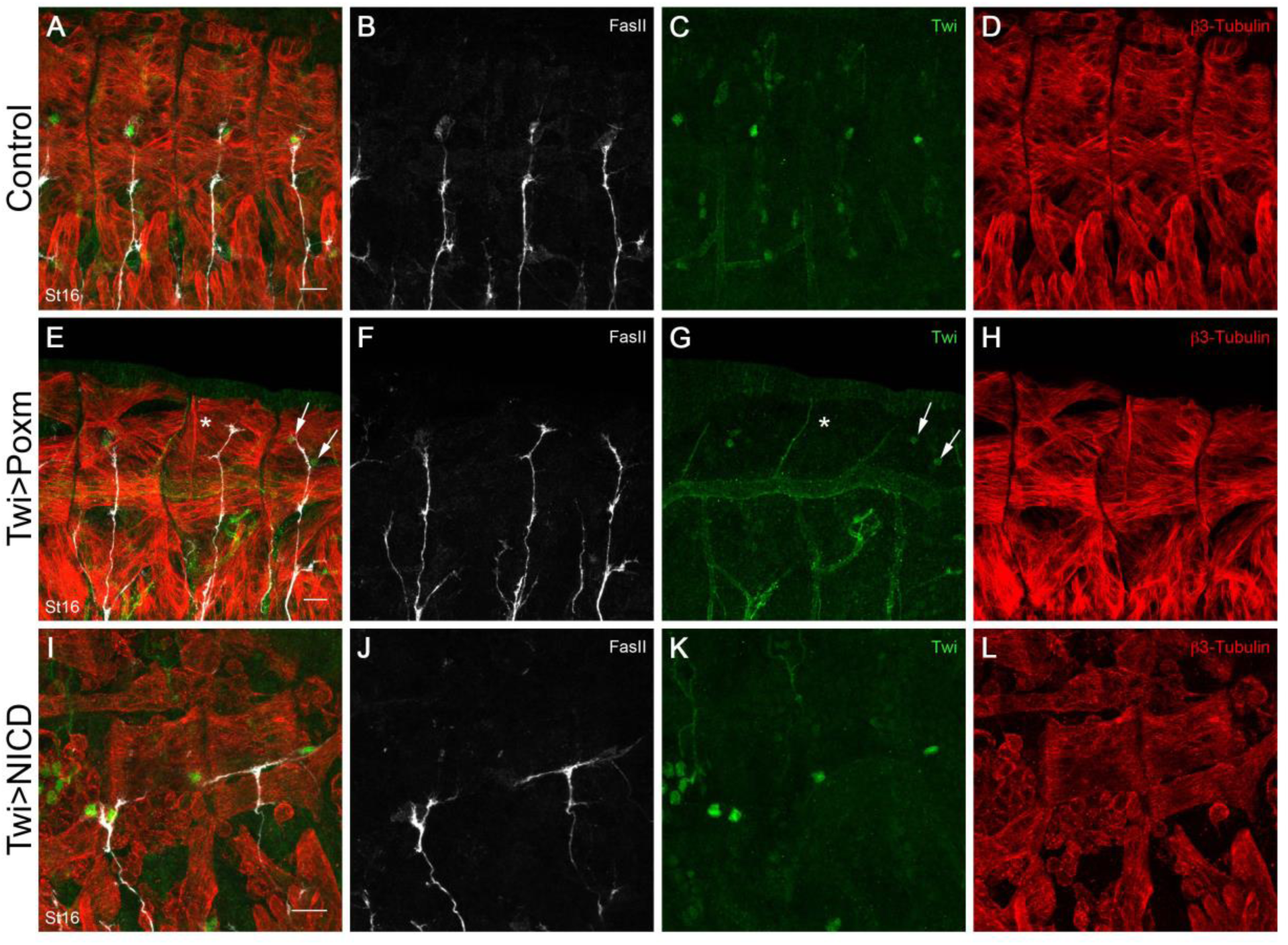
ISN is attracted by D-AMPs but its dorsal migration is not affected by their absence. Zoomed dorsal view of Twist-GAL4 (**A-D**), Twist-GAL4>UAS-Pox meso (**E-H**) and Twist-GAL4>UAS-NICD (**I-L**) stage 16 embryos. (**E-H**) The ectopic expression of Pox meso can induce either the loss of D-AMPs (white asterisk) or their duplication in a given segment (white arrows). The absence of the D-AMPs does not lead to a stall phenotype for the ISN, which can still innervate the most dorsal muscles. In the segment with duplicated D-AMPs, both retain the ability to attract the ISN. (**I-L**) Pan-mesodermal expression of the Notch intra-cellular domain (NICD) leading to the stochastic disruption of most of the body-wall muscles. Surviving D-AMPs are often mislocalized inside their respective hemisegments, and are associated with altered ISN trajectory. Scale bars represent 10 µm.

## References

1. Carmena, A., Bate, M., and Jiménez, F. (1995). Lethal of scute, a proneural gene, participates in the specification of muscle progenitors during Drosophila embryogenesis. Genes Dev. 9, 2373–2383.

2. Bate, M., Rushton, E., and Currie, D.A. (1991). Cells with persistent twist expression are the embryonic precursors of adult muscles in Drosophila. Development 113, 79–89.

3. Lai, E.C., Bodner, R., and Posakony, J.W. (2000). The enhancer of split complex of Drosophila includes four Notch-regulated members of the bearded gene family. Development 127, 3441–3455.

4. Figeac, N., Jagla, T., Aradhya, R., Da Ponte, J.P., and Jagla, K. (2010). Drosophila adult muscle precursors form a network of interconnected cells and are specified by the rhomboid-triggered EGF pathway. Development 137, 1965–1973.

5. Broadie, K.S., and Bate, M. (1991). The development of adult muscles in Drosophila: ablation of identified muscle precursor cells. Development 113, 103–118.

6. Aradhya, R., Zmojdzian, M., Da Ponte, J.P., and Jagla, K. (2015). Muscle niche-driven Insulin-Notch-Myc cascade reactivates dormant Adult Muscle Precursors in Drosophila. Elife 4.

7. Vactor, D.V., Sink, H., Fambrough, D., Tsoo, R., and Goodman, C.S. (1993). Genes that control neuromuscular specificity in Drosophila. Cell 73, 1137–1153.

8. Broadie, K., and Bate, M. (1993). Muscle development is independent of innervation during Drosophila embryogenesis. Development 119, 533–543.

9. Landgraf, M., Baylies, M., and Bate, M. (1999). Muscle founder cells regulate defasciculation and targeting of motor axons in the Drosophila embryo. Curr. Biol. 9, 589–592.

10. Ruiz Gómez, M., and Bate, M. (1997). Segregation of myogenic lineages in Drosophila requires numb. Development 124, 4857–4866.

11. Duan, H., Zhang, C., Chen, J., Sink, H., Frei, E., and Noll, M. (2007). A key role of Pox meso in somatic myogenesis of Drosophila. Development 134, 3985–3997.

12. Nose, A., Mahajan, V.B., and Goodman, C.S. (1992). Connectin: a homophilic cell adhesion molecule expressed on a subset of muscles and the motoneurons that innervate them in Drosophila. Cell 70, 553–567.

13. Nose, A., Umeda, T., and Takeichi, M. (1997). Neuromuscular target recognition by a homophilic interaction of connectin cell adhesion molecules in Drosophila. Development 124, 1433–1441.

14. Meadows, L.A., Gell, D., Broadie, K., Gould, A.P., and White, R.A. (1994). The cell adhesion molecule, connectin, and the development of the Drosophila neuromuscular system. J. Cell. Sci. 107 (Pt 1), 321–328.

15. Fambrough, D., and Goodman, C.S. (1996). The Drosophila beaten path gene encodes a novel secreted protein that regulates defasciculation at motor axon choice points. Cell 87, 1049–1058.

16. Sink, H., Rehm, E.J., Richstone, L., Bulls, Y.M., and Goodman, C.S. (2001). sidestep encodes a target-derived attractant essential for motor axon guidance in Drosophila. Cell 105, 57–67.

17. Siebert, M., Banovic, D., Goellner, B., and Aberle, H. (2009). Drosophila motor axons recognize and follow a Sidestep-labeled substrate pathway to reach their target fields. Genes Dev. 23, 1052–1062.

18. Özkan, E., Carrillo, R.A., Eastman, C.L., Weiszmann, R., Waghray, D., Johnson, K.G., Zinn, K., Celniker, S.E., and Garcia, K.C. (2013). An extracellular interactome of immunoglobulin and LRR proteins reveals receptor-ligand networks. Cell 154, 228–239.

19. Li, H., Watson, A., Olechwier, A., Anaya, M., Sorooshyari, S.K., Harnett, D.P., Lee, H.-K.P., Vielmetter, J., Fares, M.A., Garcia, K.C., et al. (2017). Deconstruction of the beaten Path-Sidestep interaction network provides insights into neuromuscular system development. Elife 6.

20. Liu, W., Wei-LaPierre, L., Klose, A., Dirksen, R.T., and Chakkalakal, J.V. (2015). Inducible depletion of adult skeletal muscle stem cells impairs the regeneration of neuromuscular junctions. eLife 4. Available at: https://elifesciences.org/articles/09221 [Accessed March 1, 2019].

21. Liu, W., Klose, A., Forman, S., Paris, N.D., Wei-LaPierre, L., Cortés-Lopéz, M., Tan, A., Flaherty, M., Miura, P., Dirksen, R.T., et al. (2017). Loss of adult skeletal muscle stem cells drives age-related neuromuscular junction degeneration. eLife 6. Available at: https://elifesciences.org/articles/26464 [Accessed March 1, 2019].

22. Brand, A.H., and Perrimon, N. (1993). Targeted gene expression as a means of altering cell fates and generating dominant phenotypes. Development 118, 401–415.

23. Lavergne, G., Soler, C., Zmojdzian, M., and Jagla, K. (2017). Characterization of Drosophila Muscle Stem Cell-Like Adult Muscle Precursors. Methods Mol. Biol. 1556, 103–116.

